# A European whitefish linkage map and its implications for understanding genome-wide synteny between salmonids following whole genome duplication

**DOI:** 10.1101/310136

**Authors:** Rishi De-Kayne, Philine G.D. Feulner

**Affiliations:** Department of Fish Ecology and Evolution, Centre of Ecology, Evolution and Biogeochemistry, EAWAG Swiss Federal Institute of Aquatic Science and Technology, Switzerland; Division of Aquatic Ecology and Evolution, Institute of Ecology and Evolution, University of Bern, Switzerland

## Abstract

Genomic datasets continue to increase in size and ease of production for a wider selection of species including non-model organisms. For many of these species highly contiguous and well-annotated genomes are unavailable due to their prohibitive complexity and cost. As a result, a common starting point for genomic work in non-model species is the production of a linkage map, which involves the grouping and relative ordering of genetic markers along the genome. Dense linkage maps facilitate the analysis of genomic data in a variety of ways, from broad scale observations regarding genome structure e.g. chromosome number and type or sex-related structural differences, to fine scale patterns e.g. recombination rate variation and co-localisation of differentiated regions. Here we present both a sex-averaged and sex-specific linkage maps for *Coregonus sp. “Albock”* containing 5395 single nucleotide polymorphism (SNP) loci across 40 linkage groups to facilitate future investigation into the genomic basis of whitefish adaptation and speciation. The map was produced using restriction-site associated digestion (RAD) sequencing data from two wild-caught parents and 156 F1 offspring in Lep-MAP3. We discuss the differences between our sex-avagerated and sex-specific maps and identify synteny between *C. sp. “Albock”* linkage groups and the Atlantic salmon (*Salmo salar*) genome. Our synteny analysis confirms that many patterns of homology observed between Atlantic salmon and *Oncorhynchus* and *Salvelinus* species are also shared by members of the Coregoninae subfamily.

## Introduction

Although advances in sequencing technology continue to increase the yield and lower the cost of genomic data acquisition, the curation of this data into a usable format can still be challenging (Ellegren 2014). Understanding the relative positions of genetic markers is often essential for the detailed analysis of genomic datasets, and is carried out in many model organisms by mapping reads to a reference genome (Sarropoulou 2011; Wolf and Ellegren 2017). However, marker ordering in the absence of a reference genome can also be carried out using a linkage map, which provides a measure of recombination distance rather than a physical difference, and as a result their production has become a common early step in the analysis of large genomic datasets (Lander and Green 1987; Lander and Schork 1994; Gross *et al*. 2008). Linkage maps are produced by observing recombination events which have occured in parents by sequencing many offspring of that parental cross. Recombination events, which break up parental combinations of alleles, are used to assign markers to, and then order within, linkage groups, elucidating the relative location of thousands of markers along the genome (Sturtevant 1913; Rastas *et al*. 2013). The resulting maps hold information on the broad genome structure e.g. number and length of linkage groups (i.e. chromosomes) and can be used to evaluate synteny with related taxa to investigate genome evolution (Sarropoulou 2011; Hale *et al*. 2017; Leitwein *et al*. 2017). Linkage maps can be used to associate phenotypes and genotypes through quantitative trait locus (QTL) mapping (Doerge 2002). Linkage maps also hold the information to investigate the colocalization of regions under selection e.g. F_ST_ outliers identified from genome scans and the recombination landscape itself (Sakamoto *et al*. 2000; Johnston *et al*. 2017). Empirical evidence has shown recombination to vary between species, populations, sexes and even individuals, highlighting the importance of its investigation in existing and new study organisms (Smukowski and Noor 2011; Kawakami *et al*. 2014; Stapley *et al*. 2017).

Linkage maps have become an essential tool in investigating evolution in non-model systems where the use of existing reference genomes is limited and the assembly of new *de novo* genomes is neither logistically nor financially feasible (Ellegren 2013; da Fonseca *et al*. 2016; Sutherland *et al*. 2016; Kubota *et al*. 2017; Sun *et al*. 2017; Zhigunov *et al*. 2017; Matz 2018). Many non-model organisms have specific ecological and evolutionary characteristics which make them particularly interesting for asking targeted evolutionary questions (Matz 2018). These features can include high speciation rate, remarkable numbers of species living in sympatry, high phenotypic and genomic diversity within or between populations, and unique ecological characteristics (Garvin *et al*. 2010; Ekblom and Galindo 2011; Hornett and Wheat 2012; Matz 2018). Carrying out studies to understanding the genomic basis of these phenomena relies upon the development of new primary genomic resources in these non-model systems (Matz 2018). Linkage maps are therefore an ideal starting point to study evolution in new systems and open the door for the future production of more complex genomic resources including *de novo* genomes. Scaffolds produced during *de novo* genome assembly can be anchored to a linkage map, improving the contiguity and accuracy of the assembly (Fierst 2015; Lien *et al*. 2016; Feulner *et al*. 2018).

Salmonids are a particularly interesting family of teleost fishes in terms of their ecology and evolution, having colonised and adapted to a huge range of freshwater habitats (Nelson *et al*. 2006). They also have an interesting evolutionary history, influenced by a whole genome duplication which occurred ~80 Mya in the shared ancestor of all salmonids (Macqueen and Johnston 2014). The family Salmonidae comprises of two main clades, which diverged ~52 Mya (Macqueen and Johnston 2014). One clade is made up of the subfamily Salmoninae which includes salmon, trout and char species and the other contains the two subfamilies Thymallinae, containing grayling, and Coregoninae, containing whitefish and ciscos (Near *et al*. 2012; Macqueen and Johnston 2014). Whitefish exhibit remarkable phenotypic diversity and high speciation rate, with multiple sympatric species having evolved post-glaciation in the last 15000 years (Lu and Bernatchez 1999; Kottelat and Freyhof 2007; Hudson *et al*. 2011). Two main whitefish species complexes exist, one in North America and the other in Europe. The North American whitefish complex comprises of *C. clupeaformis* species including sympatric ‘dwarf’ and ‘normal’ morphs which have arisen since the last glacial maximum (Bernatchez and Dodson 1990). The European species complex was previously described under the umbrella term ‘C. *laveratus* species complex’, however ongoing work to formally describe the many species which are found across Europe is being undertaken by taxonomists (Douglas *et al*. 1999; Østbye *et al*. 2005; Kottelat and Freyhof 2007; Hudson *et al*. 2011). In Europe, whitefish are naturally found as far north as Finland and as far south as the Alps, with a particularly speciose monophyletic clade known as the ‘pre-alpine’ whitefish which are distributed throughout Switzerland and its surrounding countries (Østbye *et al*. 2005; Hudson *et al*. 2011). Over 30 whitefish species have been described in Switzerland alone (Steinmann 1950) and some lakes continue to harbour up to six sympatric whitefish species despite the reduction of genetic and phenotypic differences between many species and the extinction of others following lake eutrophication in the 1980s (Vonlanthen *et al*. 2012). Sympatric whitefish species are monophyletic in many Swiss lakes and occupy a variety of ecological niches and exhibit a range of morphological differences (including body size, gill raker number and spawning season and depth; Douglas *et al*. 1999; Hudson *et al*. 2011; Vonlanthen *et al*. 2012; Hudson *et al*. 2017). It is the repeated ecological differentiation in sympatry that makes Swiss whitefish a particularly interesting radiation in which to study the genomic basis of adaptation. Although multiple studies have investigated the genetic basis of adaptation in other salmonids those carried out on the Coregoninae subfamily are comparatively scarce.

The complex evolutionary history of salmonids, specifically the effect of the salmonid specific whole genome duplication (Ss4R), makes the genetic basis of adaptation difficult to study in this family (Lien *et al*. 2016). Dense linkage maps have been produced to address these difficulties for a variety of Salmoninae, including Arctic char, Brook trout, Brown trout and Chinook salmon (McKinney *et al*. 2016; Sutherland *et al*. 2016; Leitwein *et al*. 2017; Nugent *et al*. 2017). These studies typically pair the use of dense linkage maps with the Atlantic salmon (*Salmo salar*) reference genome to improve the genomic resolution of their analyses. However, due to the ~50 million year divergence time between Salmoninae and Coregoninae, and the limited number and density of whitefish linkage maps, the current analysis of genomic whitefish datasets is complex and limited (Rogers *et al*. 2001; Rogers and Bernatchez 2004; Rogers and Bernatchez 2007; Gagnaire *et al*. 2013). Only one whitefish linkage map produced using a restriction-site associated digestion (RAD) sequencing approach is available and was produced using data from North American whitefish (*C. clupeaformis;* Gagnaire *et al*. 2013). It includes 3438 single nucleotide polymorphism (SNP) markers resolved into 40 linkage groups (matching the karyotype of *C. clupeaformis;* Phillips and Ráb 2007) and was successfully used to investigate expression QTLs in *C. clupeaformis* (Gagnaire *et al*. 2013). However, studies which later described synteny patterns between salmonid genomes struggled to confidently resolve the relationships between lake whitefish linkage groups and other salmonid chromosomes using this map (Sutherland *et al*. 2016). The use of this map for investigating the remarkable European adaptive radiation of whitefish is also limited, due to the specificity of RAD markers and lack of knowledge about genetic differentiation between *C. clupeaformis* and European whitefish (*C. laveratus* spp. complex) (Østbye *et al*. 2005; Hudson *et al*. 2011). The production of a European whitefish linkage map is therefore essential to study evolution within these extraordinary radiations.

In this study we produce a detailed linkage map for pre-alpine whitefish using a RAD sequencing approach. We produced both sex-specific and sex-averaged linkage maps for *Coregonus sp. “Albock”*, one member of the pre-alpine whitefish clade, from one F1 lab-bred cross. Here, we describe the sex-averaged and sex-specific linkage maps of *C. sp “Albock”* and use our sex-averaged linkage map to identify synteny between *C. sp. “Albock”* and Atlantic salmon (*Salmo salar*). We identify rearrangements present between the two species which reflect the occurrence of fission and fusion events following the Ss4R whole genome duplication, some of which were confidently identified to be shared only between members of the Salmoninae subfamily in past studies. This *Coregonus* linkage map will facilitate future research regarding the genomic basis of adaptation in the adaptive radiation of Swiss whitefish as well as assisting in the *de novo* assembly of the whitefish genome.

## Materials and Methods

### Experimental cross

One F1 family comprising of two parents and 156 offspring was used for linkage map construction. Both parent whitefish were sexually ripe, adult, *Coregonus sp. “Albock”*, a formally undescribed species, which likely originates from an introduction of whitefish from Lake Constance and for which taxonomic description is in progress. The parental whitefish collected from Lake Thun in December 2016 and were crossed in vitro by mixing sperm and eggs (obtained from the cantonal hatchery) together before adding cold water to harden successfully fertilized eggs. Fertilized eggs were then placed in a flow-through system which ran 5°C lake water over the eggs for 11 weeks until they began to hatch. Before larvae had fully utilized their yolk sac they were sedated and euthanized with MS222 (50 mg/l for sedation; 200 mg/l for euthanization; buffered with sodium bicarbonate 500 mg/l) and preserved in 100% ethanol (February 2017; Animal Permit number LU03/15).

### DNA extraction, library preparation and sequencing

DNA for both parental whitefish was extracted from muscle tissue. Progeny DNA was extracted following the digestion of 176 whole larvae. Both parent and progeny DNA was extracted using QIAGEN DNeasy Blood and Tissue extraction kit. The DNA concentration of each extract was measured using the Qubit 1.0 Fluorometer. In total five RAD libraries were constructed, one comprising only of the two parental individuals and the other four comprising of 44 offspring each, following the protocol of Baird *et al*. (2008) with slight modifications. Equal amounts of DNA from each offspring (1 μg) were pooled prior to restriction enzyme digestion. Since the parental library contained only two individuals, to achieve higher sequencing depth, 18 μg DNA from each parent was used for the digestion. The restriction enzyme digestion was carried out using the *Sbf-1* enzyme, which has been shown to digest salmonid DNA effectively (Gonen *et al*. 2014; Gagnaire *et al*. 2013; Sutherland *et al*. 2016), before the digested genomic DNA was ligated to the first Illumina adaptor and individual-specific barcodes. Size selection then took place using a SageELF to retain only DNA fragments between 300 and 700 base pairs. These fragments were then amplified in a PCR after the ligation of the second Illumina adaptor. Each library was spiked with PhiX DNA (~10% of reads) before being single-strand sequenced, each on a single lane of Illumina HiSeq 2500 with 100 cycles at the Lausanne Genomic Technologies Facility (Switzerland).

### Sequence processing and genotyping

The first step of processing the sequenced reads was to remove all PhiX reads using a Bowtie2 mapping approach (using default parameters except for the number of allowed mismatches which we set to 1; Langmead and Salzberg 2012). Next, all reads from the parental library were filtered for quality using TRIMMOMATIC v.0.35 (Bolger *et al*. 2014). Bases were trimmed from the beginning and end of reads if they were below quality 3, a sliding-window approach was used with a 4 base wide window to trim bases below a quality score of 15. Reads were only retained if they had an average quality of 30 and if they were still less than 50 base pairs (bp) in length. Reads from the parental library and four offspring libraries were then demultiplexed and offspring reads were trimmed to 90 bp using the *process_radtags* module in Stacks (Catchen *et al*. 2013). Next, 20 offspring with < 1 million reads were discarded to leave both parents and 156 F1 offspring for analysis. A *de novo* reference assembly was produced by combining only reads from both parents, running the *ustacks* module in Stacks (Catchen *et al*. 2013) to identify putative SNP loci present in the parents of the cross (with a minimum coverage depth of 20) and the concatenation of these consensus stacks (Catchen *et al*. 2013). An index of this reference was then produced with Bowtie2 (Langmead and Salzberg 2012). Both parental and all offspring fasta files were aligned to the parental *de novo* reference assembly using Bowtie2 (using default parameters except for the number of allowed mismatches which we set to 1) resulting in individual alignment files. The GATK *Haplotype Caller* (Poplin *et al*. 2017) was used to call genotypes, producing a VCF file retaining only SNPs genotyped with a minimum base quality score of 20 and a minimum confidence threshold of 20, i.e. p-error 0.01. This genotype file was further filtered with VCFtools (Danecek *et al*. 2011) to leave 20635 biallelic SNPs with a minimum phred quality score of 30 with indels removed. Since only one generation of offspring are included in an F1 linkage map, the most informative loci are those that are heterozygous in one parent and homozygous in the other (e.g. maternal Aa, paternal aa or maternal aa, paternal Aa). Offspring can therefore be heterozygous or homozygous (e.g. Aa or aa in an expected ratio of 1:1) and the phasing/origin of each allele is known. In addition to these highly informative loci, loci for which both parents are heterozygous can also provide information in the offspring in certain linkage mapping programs (e.g. maternal Aa, paternal Aa). In these cases, three offspring genotypes may be observed e.g. AA, Aa, aa in an expected ratio of 1:2:1 with only homozygous offspring being informative since we know that one copy of each allele is from each parent (e.g. AA offspring or aa offspring have received one A from each parent or one a from each parent, respectively). Heterozygous offspring genotypes are uninformative since the origin of each allele is unknown (e.g. Aa offspring may have received A or a from either parent). Loci were then filtered in R leaving only informative loci segregating in these two ways as well as removing any loci with missing data in either parent. All SNPs from RAD loci with more than three SNPs were removed and one SNP was chosen at random from those RAD loci with two SNPs. Remaining loci with over 20% missing data were also removed, leaving 9757 loci for linkage mapping (R Core Team 2017).

### Linkage mapping

Linkage Map construction was carried out using Lep-MAP3 (Rastas 2017). First custom R and python scripts were used to convert the VCF file containing informative loci to Lep-MAP3 format before it was converted to a posterior probability table using the script linkage2post.awk and the *Transpose* module (Lep-MAP2; Rastas *et al*. 2016). Next Lep-MAP3 modules were used starting with the *ParentCall2* module identifying 7800 informative markers. The *Filtering2* module was then used to remove markers with significant segregation distortion (dataTolerance=0.001). Linkage groups were then identified using *SeparateChromosomes2* with a logarithm of odds (LOD) score of 16 (lodLimit = 16) and the minimum number of markers per linkage group set to 25, resolving 40 linkage groups (corresponding to the 40 whitefish chromosomes identified by karyotyping; Phillips and Ráb 2007) containing 5395 loci before within-group ordering of markers was carried out (Rastas 2017). Due to the slight stochastic variation in marker distances between runs, the *OrderMarkers2* module was used, specifying a sex-specific map (sexAveraged=0), three times on each linkage group to produce a male and a female linkage map. This procedure was then repeated specifying a sex-averaged map (sexAveraged=1). The marker orders with the highest likelihoods for each linkage group for each type of map were combined to produce the final most likely male and female sex-specific maps and one final sex-averaged map, each positioning the same 5395 SNP markers. A custom R script was used to calculate differences in the marker densities and lengths between maps and the sex-averaged map was plotted using MapChart (Voorrips 2002; R Core Team 2017).

### Synteny analysis

To identify synteny between the 29 salmon chromosomes and the 40 whitefish linkage groups, the RAD loci sequenced from the two parents of the cross, which were used for *de novo* reference production, were mapped to the *Salmo salar* genome using Stampy v. 1.0.22 (Lunter and Goodson 2010) to produce an alignment file for all reference reads. Since whitefish and Atlantic salmon are ~ 60 million years divergent and transcript analysis has shown them be 93% similar, a divergence percentage of 7% (substitutionrate=0.07) was specified during read mapping (Koop *et al*. 2008). A custom R script was then used to match the 5395 SNP loci within the complete sex-averaged map to the corresponding loci in the reference whitefish - Atlantic salmon alignment file, extracting the salmon chromosome, base pair position and mapping quality. Mapped loci were then stringently filtered by their mapping quality score (MAPQ > 30) and the salmon chromosome with the most hits was noted. Linkage groups were then ordered to reflect their synteny with salmon chromosomes (Table 1) and renamed with the prefix ‘W’ to match salmon chromosome ordering. Synteny was visualized out using the *circlize* package (Gu 2014) in R plotting all links from reads with MAPQ > 30 to the corresponding salmon chromosome and position within this chromosome (Figure 2).

**Table 1:** Table comparing statistics for the sex-averaged, female and male *C. sp. “Albock”* linkage maps. The results of synteny analysis are included, showing the homologous salmon chromosome for each whitefish linkage group and the re-ordered whitefish linkage group name.

| Linkage Group | Number of SNPs | LG length (cM) | SNPs/cM | Female LG length (cM) | Female SNPs/cM | Male LG length (cM) | Male SNPs/cM | Homologous Salmon Chromosome | Synteny LG | LG Length Sex Ratio (F:M) |
| --- | --- | --- | --- | --- | --- | --- | --- | --- | --- | --- |
| Calb01 | 253 | 75.96 | 0.30 | 91.07 | 0.36 | 63.67 | 0.25 | Ssa01 | W02 | 1.43 |
| Calb02 | 228 | 83.57 | 0.37 | 101.33 | 0.44 | 69.58 | 0.31 | Ssa01 | W03 | 1.46 |
| Calb03 | 220 | 78.51 | 0.36 | 84.40 | 0.38 | 87.95 | 0.40 | Ssa21 | W32 | 0.96 |
| Calb04 | 214 | 58.45 | 0.27 | 66.69 | 0.31 | 50.05 | 0.23 | Ssa10 | W15 | 1.33 |
| Calb05 | 190 | 66.93 | 0.35 | 63.63 | 0.33 | 71.66 | 0.38 | Ssa12 | W18 | 0.89 |
| Calb06 | 187 | 53.16 | 0.28 | 70.69 | 0.38 | 37.88 | 0.20 | Ssa13 | W20 | 1.87 |
| Calb07 | 181 | 71.53 | 0.40 | 68.13 | 0.38 | 88.06 | 0.49 | Ssa04 | W06 | 0.77 |
| Calb08 | 173 | 52.28 | 0.30 | 56.37 | 0.33 | 45.30 | 0.26 | Ssa10 | W14 | 1.24 |
| Calb09 | 170 | 79.41 | 0.47 | 73.03 | 0.43 | 91.75 | 0.54 | Ssa07 | W10 | 0.80 |
| Calb10 | 165 | 62.43 | 0.38 | 60.45 | 0.37 | 65.05 | 0.39 | Ssa01 | W01 | 0.93 |
| Calb11 | 164 | 65.01 | 0.40 | 64.04 | 0.39 | 66.05 | 0.40 | Ssa11 | W16 | 0.97 |
| Calb12 | 164 | 51.09 | 0.31 | 70.15 | 0.43 | 30.22 | 0.18 | Ssa22 | W33 | 2.32 |
| Calb13 | 162 | 69.34 | 0.43 | 71.26 | 0.44 | 63.49 | 0.39 | Ssa29 | W40 | 1.12 |
| Calb14 | 157 | 65.11 | 0.41 | 61.78 | 0.39 | 72.14 | 0.46 | Ssa13 | W19 | 0.86 |
| Calb15 | 156 | 64.90 | 0.42 | 63.19 | 0.41 | 71.73 | 0.46 | Ssa16 | W24 | 0.88 |
| Calb16 | 154 | 56.17 | 0.36 | 55.30 | 0.36 | 65.75 | 0.43 | Ssa20 | W31 | 0.84 |
| Calb17 | 151 | 65.53 | 0.43 | 69.40 | 0.46 | 61.63 | 0.41 | Ssa23 | W34 | 1.13 |
| Calb18 | 149 | 61.50 | 0.41 | 65.22 | 0.44 | 62.38 | 0.42 | Ssa09 | W11 | 1.05 |
| Calb19 | 147 | 62.15 | 0.42 | 68.25 | 0.46 | 55.50 | 0.38 | Ssa14 | W21 | 1.23 |
| Calb20 | 144 | 66.36 | 0.46 | 79.08 | 0.55 | 56.52 | 0.39 | Ssa27 | W37 | 1.40 |
| Calb21 | 143 | 71.78 | 0.50 | 69.37 | 0.49 | 83.01 | 0.58 | Ssa25 | W36 | 0.84 |
| Calb22 | 137 | 71.12 | 0.52 | 74.56 | 0.54 | 67.96 | 0.50 | Ssa03 | W04 | 1.10 |
| Calb23 | 127 | 64.80 | 0.51 | 68.96 | 0.54 | 69.78 | 0.55 | Ssa06 | W09 | 0.99 |
| Calb24 | 127 | 52.57 | 0.41 | 58.54 | 0.46 | 54.23 | 0.43 | Ssa15 | W22 | 1.08 |
| Calb25 | 124 | 57.74 | 0.47 | 61.62 | 0.50 | 60.81 | 0.49 | Ssa24 | W35 | 1.01 |
| Calb26 | 123 | 64.59 | 0.53 | 70.67 | 0.57 | 62.12 | 0.51 | Ssa19 | W29 | 1.14 |
| Calb27 | 118 | 46.03 | 0.39 | 61.06 | 0.52 | 30.24 | 0.26 | Ssa18 | W27 | 2.02 |
| Calb28 | 115 | 59.05 | 0.51 | 63.68 | 0.55 | 59.73 | 0.52 | Ssa15 | W23 | 1.07 |
| Calb29 | 114 | 62.40 | 0.55 | 61.31 | 0.54 | 70.58 | 0.62 | Ssa09 | W12 | 0.87 |
| Calb30 | 112 | 62.75 | 0.56 | 68.12 | 0.61 | 63.96 | 0.57 | Ssa05 | W08 | 1.07 |
| Calb31 | 111 | 53.35 | 0.48 | 63.62 | 0.57 | 42.48 | 0.38 | Ssa20 | W30 | 1.50 |
| Calb32 | 104 | 56.67 | 0.54 | 63.47 | 0.61 | 53.94 | 0.52 | Ssa18 | W28 | 1.18 |
| Calb33 | 97 | 67.73 | 0.70 | 70.46 | 0.73 | 66.40 | 0.68 | Ssa09 | W13 | 1.06 |
| Calb34 | 79 | 61.12 | 0.77 | 71.34 | 0.90 | 62.97 | 0.80 | Ssa03 | W05 | 1.13 |
| Calb35 | 56 | 36.88 | 0.66 | 55.57 | 0.99 | 21.14 | 0.38 | Ssa28 | W38 | 2.63 |
| Calb36 | 45 | 24.18 | 0.54 | 15.92 | 0.35 | 30.75 | 0.68 | Ssa17 | W26 | 0.52 |
| Calb37 | 37 | 27.48 | 0.74 | 34.82 | 0.94 | 21.51 | 0.58 | Ssa11 | W17 | 1.62 |
| Calb38 | 34 | 11.86 | 0.35 | 0.00 | 0.00 | 24.01 | 0.71 | Ssa16 | W25 | 0.00 |
| Calb39 | 32 | 17.17 | 0.54 | 0.00 | 0.00 | 33.66 | 1.05 | Ssa04 | W07 | 0.00 |
| Calb40 | 31 | 15.20 | 0.49 | 23.55 | 0.76 | 7.41 | 0.24 | Ssa28 | W39 | 3.18 |
| Total | 5395 | 2293.86 |  | 2460.10 |  | 2263.05 |  |  |  | 1.09 |
| Average | 134.88 | 57.35 | 0.46 | 61.50 | 0.48 | 56.58 | 0.46 |  |  |  |

### Data availability

Fastq files for all 156 offspring and both parents are deposited in the NCBI short read archive (SRA accession xxxx available upon publication). The genotype file (VCF) and the Lep-MAP input file are available at Dryad (doixxx, available upon publication). All R, Python and bash scripts used can be accessed at https://github.com/RishiDeKayne/.

## Results and Discussion

### Linkage mapping

Our F1 cross comprising of two *C. sp. “Albock”* adults and 156 offspring was successfully genotyped using a RAD-seq approach. The assignment of SNPs to linkage groups and the subsequent ordering of SNPs within linkage groups was carried out using Lep-Map3 (Rastas 2017). In total 9757 SNPs were retained following stringent quality control and loci filtering steps, with 7800 identified as informative in Lep-MAP3. Finally, 5395 SNPs were assigned to, and arranged within, linkage groups in both sex-averaged and sex-specific maps (Table 1; Figure 1). With the LOD score of 16, 40 linkage groups, corresponding to the 40 chromosomes observed in karyotype studies of European whitefish (*C. laveratus;* Phillips and Ráb 2007), were formed with an average of 135 markers per linkage group (Table 1). Map lengths varied from 2293.86 cM in the sex-averaged map to 2460.10 cM and 2263.05 cM in the female and male maps, respectively. All three maps produced in this study were considerably shorter than a prevoiusly published *C. clupeaformis* linkage map containing 3438 RAD markers, which had a total map length of 3061 cM (Gagnaire *et al*. 2013). Our sex-averaged *C. sp. “Albock”* map had an average linkage group length of 57.35 cM with the female and male sex-specific maps showing average linkage group lengths of 61.50 cM and 56.58 cM, respectively.

**Figure 1:**
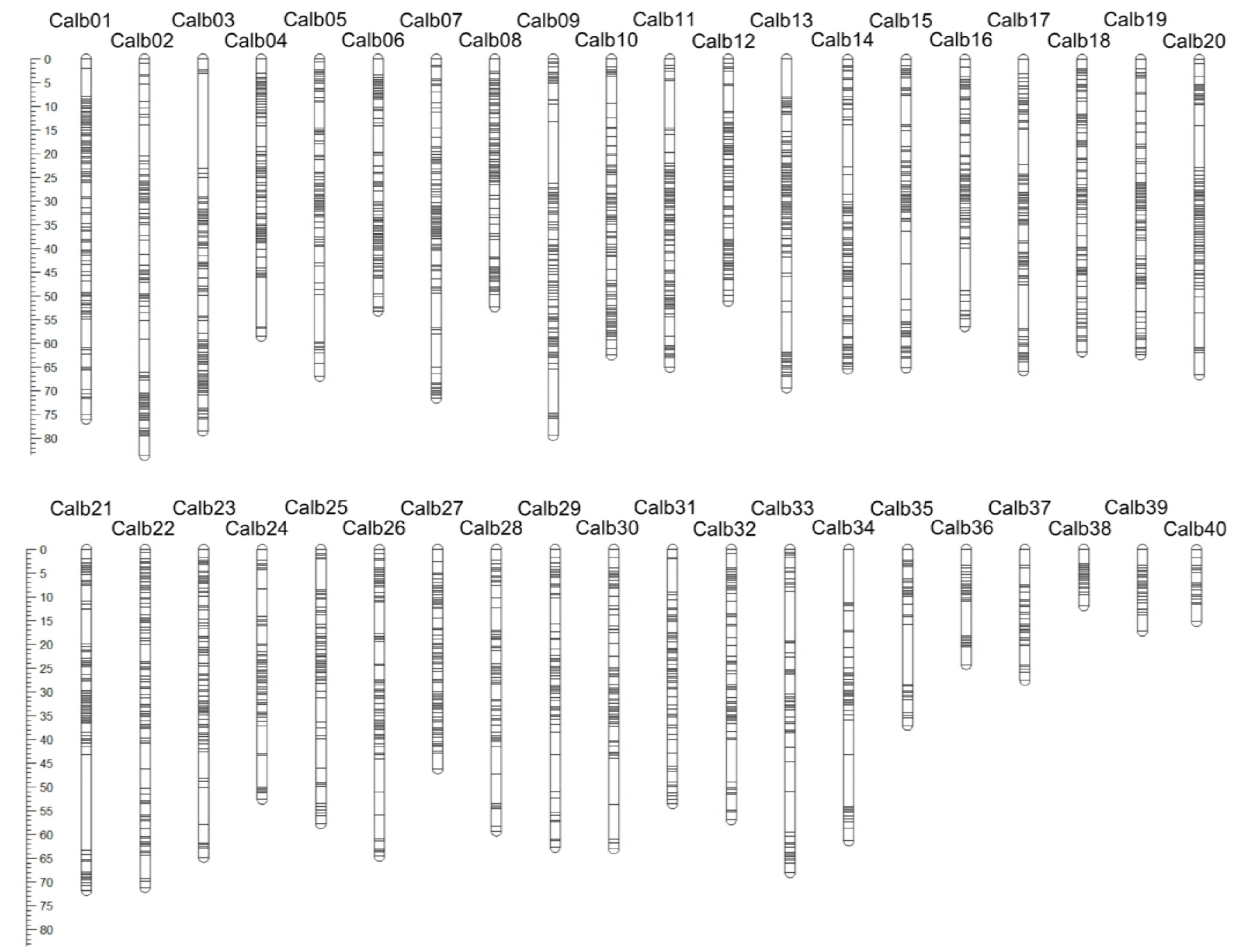
European whitefish linkage map showing the grouping and position of 5395 SNPs within a sex-averaged linkage map. The length of each of the 40 linkage groups is indicated by the scale in cM with linkage groups ordered by marker number from highest to lowest.

**Figure 2:**
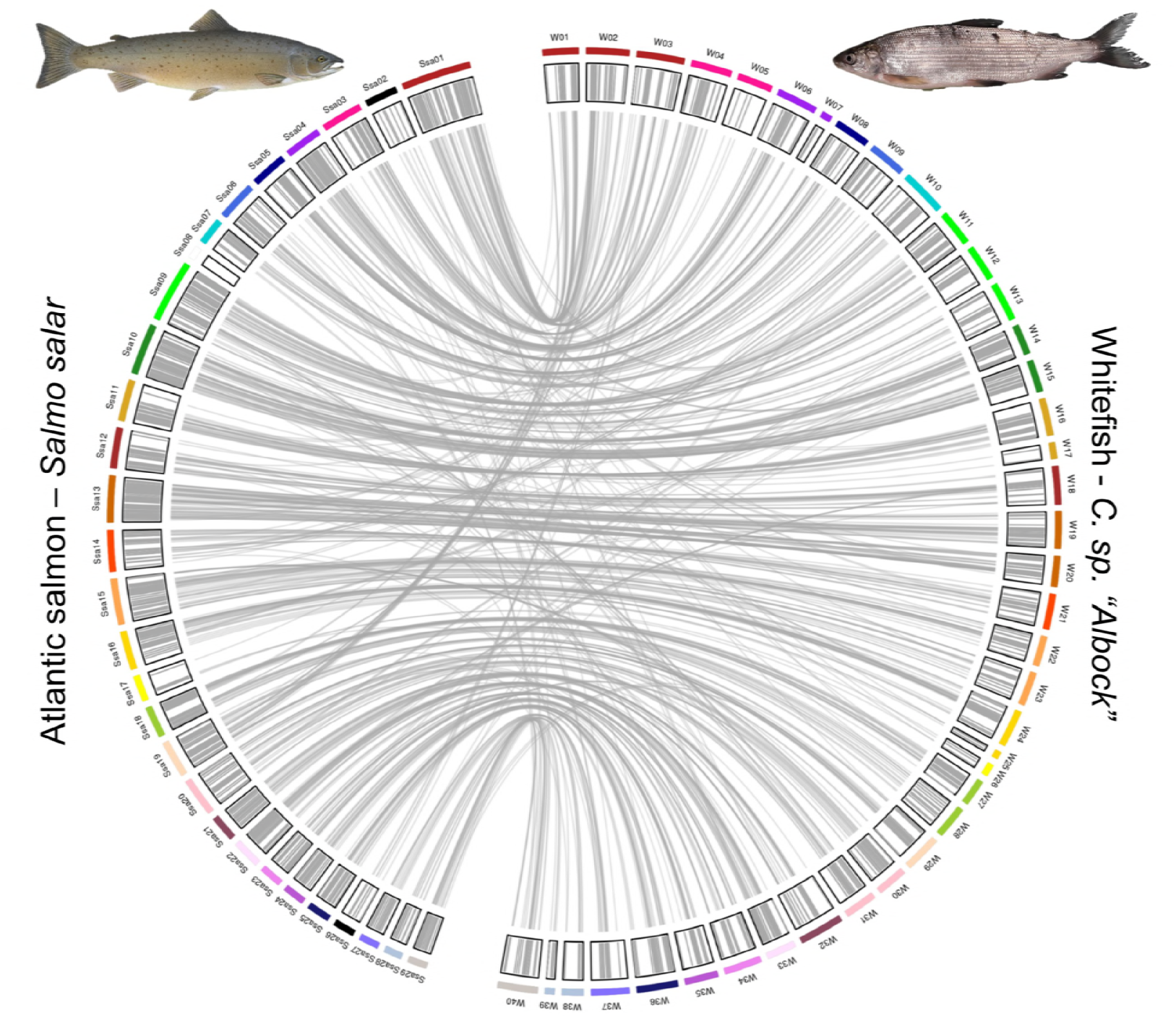
Synteny plot identifying homologous whitefish linkage groups and Atlantic salmon (*Salmo salar*) chromosomes denoted by the colouring of linkage groups and chromosomes. Links represent the location of 839 markers within the whitefish linkage map which were successfully mapped to the Atlantic salmon genome. Black salmon chromosomes Ssa02 and Ssa26 represent chromosomes with no homologous whitefish linkage groups.

The number of SNPs per linkage group varied from 31 to 253 and the lengths of linkage groups varied from 15.20 cM to 83.57 cM in the sex-averaged map. Two linkage groups, Calb38 and Calb39, were comprised only of male-informative loci and therefore had lengths of 0 cM in the female map, with the longest linkage group in the female map being Calb02 at 101.33 cM. In the male map linkage groups vary in length from 7.41 cM to 88.06 cM for linkage groups Calb40 and Calb07.

Our sex-averaged map has high resolution, with a low average cM per marker of 0.46 cM, varying from 0.27 cM in Calb04 to 0.77 cM in Calb34. The linkage map of the close relative *C. clupeaformis*, a representative of the North American whitefish lineage, had a marker resolution across the map of 0.89 cM, around half the density of our *C. sp “Albock” map*. In the female map the average cM per marker was 0.48 cM varying in linkage groups (only considering linkage groups > 0 cM) from 0.31 cM in Calb04 to 0.99 cM in Calb35. The average cM per marker in the male map was 0.46 cM with the smallest and largest ratios found in Calb12 and Calb39 respectively with 0.18 cM and 1.05 cM.

Sex differences can be observed by comparing our sex-specific linkage maps for *C. sp*. *“Albock”*. Comparing total map lengths for the female and male maps gives a sex-ratio (female:male) of 1.09 however, this does not account for the two whitefish linkage groups which have length 0 cM in our female map (Calb38 and Calb39). Calculating this length sex-ratio for each linkage group separately, including only those > 0 cM in both maps, results in a sex-ratio of 1.25. Salmonid species have been shown to have sexual dimorphisms in recombination rate with female:male map length ratios for these species varying from 1.38 in Atlantic salmon (Lien *et al*. 2011) to 2.63 in Brown trout (Gharbi *et al*. 2006) and therefore sexual dimorphism in whitefish appears to be low in comparison to other salmonids. However, since each sex-specific linkage map represents the recombination landscape in one individual, in our case each parent of the F1 cross, more than one linkage map is required to disentangle individual variation in recombination rate and consistent sex specific recombination rate variation (Sakamoto *et al*. 2000; Moen *et al*. 2004; Lien *et al*. 2011). Although our sex-ratio does not conclusively show variable recombination rates between females and males it still reveals a striking difference in map length considering the inclusion of the same set of markers for each. Studies on other teleost species, including stickleback, have reported detailed empirical evidence of sexually dimorphic recombination rates, calculating sex-ratios of linkage map lengths to be 1.64 (Sardell *et al*. 2018). Future work should aim to compare and contrast the recombination landscape of whitefish to the detailed sexually dimorphic recombination patterns observed in drosophila, mice, deer and various fish species (Dunn 1920; Sakamoto *et al*. 2000; Lenormand and Dutheil 2005; Johnston *et al*. 2017; Kubota *et al*. 2017; Sardell *et al*. 2018).

### Synteny analysis

Synteny analysis was carried out to investigate broad scale genome structural variation, such as fission and fusions of chromosomes or chromosome arms, within the Salmonidae family. Stringent filtering of mapped RAD-seq reads to the salmon genome was applied to identify synteny whilst excluding uncertain mappings. From 5395 loci included in our linkage map we retained 839 mappings of high quality, which were spread across all 40 whitefish linkage groups (Figure 2). Homology between salmon chromosomes and whitefish linkage groups was determined by identifying the most common salmon chromosome the markers on each whitefish linkage group mapped to. Only a small number of markers on each whitefish linkage group mapped to a different salmon chromosome than the identified homologous chromosome (shown by the low abundance of non-parallel links from each linkage group; Figure 2). The largest of these deviations is a series of links (16) from W02 (which was identified as homologous to Ssa01 with 18 links) to Ssa19. Due to the similar abundance of links to two different salmon chromosomes these are unlikely to be erroneous or uncertain mappings and might rather reflect a whitefish specific fusion of two Atlantic salmon chromosomes Ssa01 and Ssa19, although future research is required to confirm this. Three salmon chromosomes, Ssa02, Ssa08 and Ssa26 were not identified as homologs to any of our whitefish linkage groups, with Ssa08 having no significant mappings at all. This might be due to the fact that this salmon chromosome is small (26.43 Mb) and may also be small in whitefish, decreasing the number of RAD loci sequenced within it by chance (Lien *et al*. 2016). However, the *C. clupeaformis* linkage map also failed to identify synteny to Ssa08, raising the possibility this chromosome has been lost in whitefish (Gagnaire *et al*. 2013; Sutherland *et al*. 2016). We identified two salmon chromosomes which were each homologous to three different whitefish linkage groups; Ssa01 to W01, W02 and W03 and Ssa09 to W11, W12 and W13 (Figure 2). These Atlantic salmon chromosomes have been identified to map to three linkage groups in other salmonids including Brook Charr, Arctic Charr, Coho salmon and various *Oncorhynchus* species, however, synteny with *C. clupeaformis*, the only member of Coregoninae included in these comparisons, was less clear (Kodama *et al*. 2014; Sutherland *et al*. 2016; Hale *et al*. 2017; Nugent *et al*. 2017). This syntenic pattern has been attributed to fusion events which were unique to the Atlantic salmon lineage only. Here we add to the evidence provided by the *C. clupeaformis* linkage map that this synteny is also consistent with Coregoninae despite their significant divergence from members of the Salmoninae. Synteny analysis between members of Salmonidae also identified a number of Atlantic salmon chromosomes which each show homology to two linkage groups (Sutherland *et al*. 2016; Hale *et al*. 2017). We find a similar pattern of synteny between *Salmo salar* and *Coregonus* for many of these salmon chromosomes including Ssa03 (to W04 and W05), Ssa10 (to W14 and W15), Ssa13 (to W19 and W20), Ssa15 (to W22 and W23), Ssa16 (to W24 and W25), Ssa18 (to W27 and W28) and Ssa20 (to W30 and W31) (Figure 2). Our synteny analysis also identified Ssa04 as homologous to W06 and W07 and Ssa11 as homologous to W16 and W17, although W07 and W17 have very few links (1 and 2, respectively). We also find that the multiple one to one relationships between salmon chromosomes and salmonid linkage groups identified by Sutherland *et al*. (2016) are also consistent with our map including those to Ssa12 (W18), Ssa22 (W33), Ssa23 (W34), Ssa24 (W35), Ssa25 (W36), Ssa27 (W37) and Ssa29 (W40; Table 1). We identify one possible whitefish-specific fission event with markers from both W38 and W39 mapping to Ssa28, which is homologous to only one linkage group in each salmonid species compared by Sutherland *et al*. (2016) including *C. clupeaformis*. It is therefore possible that a fission event has occurred in the European whitefish lineage, however, due to relatively low number and density of markers on W38 and W39 future investigation should aim to clarify this pattern. Our synteny analysis also highlights possible fusion events which have occured only in European whitefish and Atlantic salmon. Whilst the *C. clupeaformis*, Rainbow trout, and multiple salmonid linkage maps, identified synteny of Ssa05, Ssa06, Ssa14, Ssa17 and Ssa19 to two linkage groups each, we only identify synteny to W08, W09, W21, W26 and W29, respectively (Table 1; Gagnaire *et al*. 2013; Sutherland *et al*. 2016). Two salmon chromosomes, Ssa07 and Ssa21 were shown by *Sutherland et al*. (2016) to be homologous to two linkage groups in *C. clupeaformis* but only one linkage group in all other salmonids. Our *C. sp*. *“Albock”* map identifies Ssa07 as homologous to W10 and Ssa21 to W32 suggesting the pattern of synteny may not be conserved between *Coregonus* species. Further work must therefore be carried out to determine genome structure similarity between *C. sp. “Albock”* and *C. clupeaformis*.

Both broad and small scale structural variations, including inversions, duplications and deletions, have been observed between closely related species and the mis-segregation which can occur during meiosis as a result of these variations is thought to be able to play a role in the speciation process (Feulner and De-Kayne 2017). It is therefore possible that European and North American whitefish lineages (and even species within these lineages) have unique structural variations which may underpin reproductive isolation in sympatry. Without information on genome wide synteny and the occurrence of structural variation between these two lineages it is difficult to determine whether the observed variation in synteny patterns to the Atlantic salmon (e.g. with regards to Ssa05, Ssa06, Ssa14, Ssa17 and Ssa19) represents true variation between these species or variation in linkage mapping resolution and accuracy. A comparison of synteny between our *C. sp. “Albock”* map and the Atlantic salmon (using our synteny mapping approach) and the *C. clupeaformis* map to the Atlantic salmon (compared by Sutherland *et al*. 2016) can be found in Table S1.

### The development of genomic resources for European whitefish

A wealth of genomic resources used to study adaptation and speciation are now available for a variety of systems. Multiple species from popular model radiations including Galapagos finches and Lake Victoria cichlids now have highly contiguous, well curated and annotated, reference genomes (Brawand *et al*. 2014; Lamichhaney *et al*. 2015). These resources provide the opportunity to ask specific questions about intra and inter-species genomic differences with many studies focusing on understanding the genomic basis of adaptation and reproductive isolation. Studies can now utilize high throughput whole-genome sequencing to achieve high depth of coverage and are able to map these reads to a reference genome to understand the distribution of genomic variation along the genome.

However, many interesting organisms including the many ecologically diverse salmonids have limited reference genomes available. Although both the Atlantic salmon (*Salmo salar*) and Rainbow trout (*Oncorhynchus mykiss*) genomes have been assembled (Berthelot *et al*. 2014; Lien *et al*. 2016), they are distantly related to many species within the salmon family, specifically the diverse clade of whitefish. Whitefish show extreme phenotypic diversity, although the degree of differentiation in sympatry varies. In North America a variety of lakes harbour sympatric pairs of ‘normal’ and ‘dwarf’ whitefish (Bernatchez and Dodson 1990). In Europe the diversity of whitefish species is much greater, and in Switzerland specifically up to six species of pre-alpine whitefish exist in sympatry in some post-glacial lakes (Hudson *et al*. 2011; Hudson *et al*. 2017). These species differ in many phenotypic traits including gill-raker number, body size and body shape. Reproductive isolation between species is thought to be largely maintained by different spawning seasons and depths between sympatric species (Hudson *et al*. 2017). Because of the rapid post-glaciation diversification of whitefish, the genomic basis of adaptation and speciation in pre-alpine whitefish is of particular interest. However, the best assembled, most closely related reference genome to this whitefish radiation is that of the Atlantic salmon. Since these two clades are so distantly related, new whitefish-specific resources are needed. Despite the production of multiple *Coregonus clupeaformis* linkage maps (Rogers *et al*. 2001; Rogers and Bernatchez 2004; Rogers and Bernatchez 2007), only one map to-date (Gagnaire *et al*. 2013) has utilized a high throughput sequencing technique. Although this map contains many SNP loci its use is largely restricted to the North American system, since the genome-wide synteny between North American and European whitefish is unknown and RAD markers are anonymous.

Our linkage map fills a gap in the resources available to analyse European whitefish genetic data allowing investigation into this species rich, ecologically diverse, lineage. The patterns of synteny between European whitefish and Atlantic salmon reported here should be further investigated once whitefish genomes become available to identify synteny at a finer scale, identifying chromosome fission and fusion events and possible inversions also within the *Coregonus* genus. Our linkage map can also be paired with future resources to investigate the outcome of whole genome duplication including estimations of the rediploidized proportion of the genome, already calculated in Atlantic salmon and Rainbow trout. Future work should further aim to identify regions of the genome which may underpin reproductive isolation in whitefish to better understand the speciation mechanism in this adaptive radiation.

In conclusion, we have produced the densest *Coregonus* linkage map to date, with a total sex-averaged map length of 2293.86 cM containing 5395 SNP loci. We have found evidence of sex-specific recombination rate variation within *C. sp. “Albock”* by calculating the sex-ratio in female and male linkage map lengths. The level of heterochiasmy inferred by this sex-ratio is reflected in other species with known sex-specific recombination variation, including other salmonids (Gharbi *et al*. 2006; Lien *et al*. 2011). We also show that *C. sp. “Albock”* linkage groups exhibit synteny with Atlantic salmon chromosomes, in some cases following a pattern of synteny shared with other salmonid species. This linkage map will facilitate a host of future studies into the genomic basis of adaptation in pre-alpine whitefish including those on the identification of QTLs for traits of interest, the interpretation of genome-wide divergence data and the colocalization of regions under selection e.g. F_ST_ outliers identified from genome scans. It also has the potential to assist in future scaffolding of pre-alpine whitefish reference genomes.

## Acknowledgements

Thanks to Benjamin Gugger and team from Lake Thun whitefish hatchery for providing us with the breeding pair of *C. sp. “Albock”*. Also thanks to Anna Feller, David Frei, Andreas Taverna and Erwin Schäffer for their help breeding and maintaining the whitefish larvae and Oliver Selz for his taxonomic expertise. This project is funded by the Swiss National Science Foundation (SNSF project 31003A_163446/1 awarded to PGDF).

---

**Table S1:**
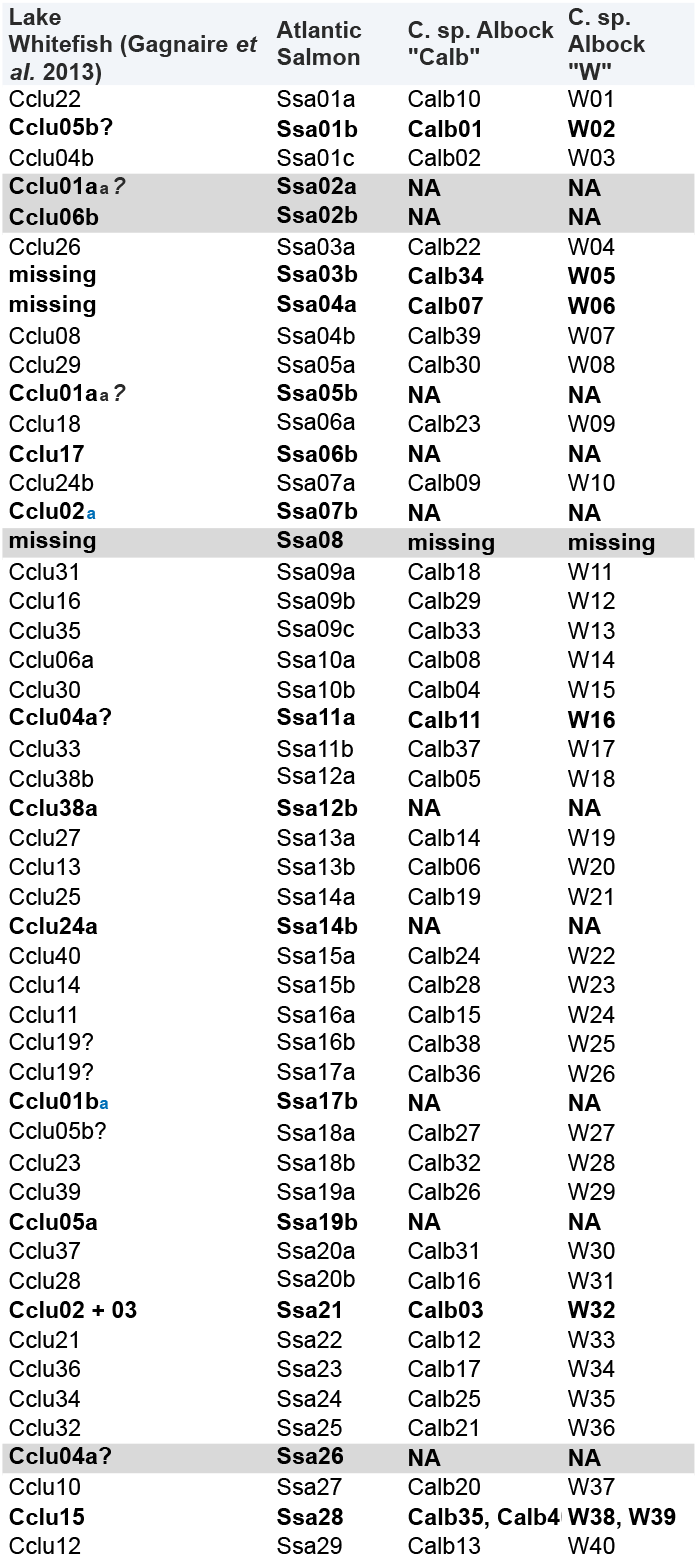
Table comparing the synteny identified between North American lake whitefish (*Coregonus clupeaformis*) and Atlantic salmon (*Salmo salar*) by Sutherland et al. (2016) using MapComp and our identified synteny between pre-alpine whitefish (*C. sp. “Albock”*) and Atlantic salmon using a mapping approach.

